# ARCHER-LD: Rapid Long-Range Linkage Disequilibrium Calculations at Biobank Scale using GPU Acceleration

**DOI:** 10.64898/2026.09.15.751759

**Authors:** Rachit Kumar, Pankhuri Singhal, David Zhang, Christopher Carson, Mitchell Conery, Alex Rodriguez, Tarak Nath Nandi, Marylyn D. Ritchie, Mathialakan Thavappiragasam, Benjamin F. Voight, Ravi K. Madduri, Anurag Verma

## Abstract

Linkage disequilibrium (LD) information from one’s own dataset is considered optimal for downstream analyses such as statistical fine-mapping. However, the computational complexity (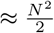 computations for N variants) lead most studies to use external reference panels, such as from 1000 Genomes. To capture LD across all variant pairs in biobank-scale whole-genome sequencing (WGS) datasets with hundreds of millions of variants, new computational strategies are essential. We present a novel approach that uses multi-GPU distributed computing to compute *R*^2^ for every variant pair in a dataset. On chromosome 22 (1.8 million variants) of the 30x WGS 1000 Genomes dataset, our method using eight consumer-level 12GB GPUs (NVIDIA RTX 2080TIs) takes <10 minutes, while the same calculation with PLINK using a high-end 64- threaded CPU (Intel Xeon Gold 6338) takes >80 minutes, corresponding to a ≈8x speedup. We further demonstrate true biobank-scale performance in the Penn Medicine Biobank (PMBB; 57,170 samples), computing chromosome 1 LD (≈1.38 million variants) in 49 minutes versus 22.7 hours for PLINK, a ≈28x speedup. With this method, we successfully computed on the aforementioned 30x WGS 1000 Genomes dataset (≈120 million variants and ≈2500 samples) the entire LD matrix (>1e16, or 10 quadrillion elements) in under 6 hours using 512 NVIDIA 40GB A100 GPUs on the Department of Energy Argonne Leadership Computing Facility Polaris Supercomputer. We make this tool, coded in Python using CuPy, publicly available. Using this tool, researchers can leverage the full extent of their genomic data without relying on external LD reference panels and acquire more accurate, population-specific findings, particularly for groups underrepresented in existing databases.

**Key Messages:** - ARCHER-LD makes use of GPUs to speed up the computation of linkage disequilibrium dramatically.
- After conversion of genotype data to a disk-based matrix, we can compute LD in parallel over chunks of the genotype matrices for both phased and unphased data.
- By making use of a parallel formulation of LD calculations, ARCHER-LD can scale to any number of GPUs that are available.
- ARCHER-LD scales to real biobanks: in the Penn Medicine Biobank (57,170 samples) it computes whole-chromosome LD substantially faster than PLINK, with the speedup increasing as the dataset grows.
- ARCHER-LD will be publicly available for use in calculating in-sample LD and extension by other researchers.

## Introduction

Linkage disequilibrium (LD) is a phenomenon in genomics where genetic variants are, in a population, associated with each other more often than expected by chance [1]. As a result, many standard methods to identify variants that can cause or contribute to disease cannot definitively pinpoint a causal variant, since any given variant may be in LD with another variant inherited alongside it.

An important step in genomics research is to compute linkage disequilibrium maps that characterize how variants associate with each other. However, computing such maps is an O(*N* ^2^) computation for N variants and scales poorly to biobank-scale datasets with millions of variants. To make the computation tractable in time, many current approaches assume that LD occurs only between variants in close proximity by imposing predefined distance thresholds when selecting pairs for LD calculation. By construction, this discards all LD between distant variants; for example, the correlation between a variant on chromosome 6 and a variant on chromosome 17 is never computed, so it cannot be observed. However, such long-range and even trans-chromosomal LD is real and biologically meaningful: it is well documented at loci such as the major histocompatibility complex (MHC) and known chromosomal inversion sites, where it can confound genome scans and ancestry inference if not identified [2, 3]. Such long-range LD can be observed for a variety of reasons that are not fully understood, with some of the most common hypotheses including nonrecombination-based mechanisms such as epistatic selection [4, 5].

Furthermore, computing linkage disequilibrium from the dataset in which you seek to apply it (also known as in-sample LD) is generally considered ideal for downstream analyses due to capturing the nuances of population structure and diversity that exist within your actual dataset, if the dataset is large enough [2, 6]. However, in-sample LD is often considered infeasible to compute due to the fact that a larger sample size means increased coverage and inclusion of rare and ultra-rare variants coupled with the aforementioned O(*N* ^2^) scaling. This leads many researchers to use reference panels where LD has been precomputed in a different population, such as the 1000 Genomes dataset, rather than computing in-sample LD on their own dataset [6, 7].

In this paper, we present Accelerated Rapid Computation of High-throughput Extensive Recombination and Linkage Disequilibrium (ARCHER-LD), a new method that leverages multiple GPUs to perform rapid genome-wide LD calculations. ARCHER-LD enables biobank-scale computations of LD from phased or unphased genotype data in a tractable and efficient manner, overcoming the limitations of traditional distance-based approaches.

We validate ARCHER-LD both on the 1000 Genomes Project and, importantly, at true biobank scale in the Penn Medicine Biobank (57,170 samples), where its speed advantage over existing tools is largest and continues to grow with the number of samples and variants.

## Methods

### Genotype Data Processing and Storage

Most large genotype data are stored in a text-based format known as the Variant Call Format (VCF), where loading genotypes from this format can be slow. To address this, we preprocess in a step wherein VCF files are converted to Zarr format using cyvcf2 [8] and Kerchunk [9]. Some genotype data are also stored in the form of PLINK binary datasets as these can be more performant; we leverage a utility known as bio2zarr to generate Zarr representations of these datasets with some additional processing performed to compute metadata for variants [10].

Zarr allows for on-disk stores of data in an efficient manner that allows for quick, chunked loading of the data from disk, essential for reducing disk-read bottlenecks for the LD matrix calculations and ensuring that the GPU steps are rate-limiting [11]. This is done on the CPU only, as file read currently cannot be sped up by the GPU. The resulting Zarr files then enable speedy, on-the-fly, chunked retrieval of the genotype data from disk, which is essential for the primary computations in ARCHER-LD without fully loading the entire genotype matrix into memory.

The genotype matrix denoted as *A* is of dimension N variants x K samples x 2 haplotypes and is stored as an int8 data type. Each element of *A* is one of 0 (reference), 1 (alternate), or -1 (missing). For phased data, *A*_*x*1_ represents the first haplotype of an arbitrary row x of the matrix, and *A*_1_, *A*_2_ each represent the individual haplotype matrices. Unphased data is stored identically, with heterozygotes having the variant always placed on the first haplotype matrix (*A*_1_). Some metadata is stored alongside the genotype matrix to enable downstream variant identification and filtering by the user.

### Computation of the Phased *R*^2^ LD Matrix

In ARCHER-LD, we compute the LD matrix for phased data, defined as a matrix denoted as *R*^2^ (where each element is a phased correlation coefficient 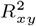 ) for all N variants (“features”) in a dataset pairwise, with *K* samples and two haplotypes for each variant. The value of *R*^2^ can be computed using a related matrix called *D* (where each element is the coefficient of linkage disequilibrium *D*_*xy*_ as described in [1]). *D* can be represented by the elementwise calculation for the genotype matrix *A*, where *A*_*xn*_ represents the corresponding variant row *x* and haplotype *n* of *A, N*_*xy*_ represents the joint haplotype sample size (of non-missing calls) of variant *x* and *y*, and *N*_*x*_ or *N*_*y*_ represent the haplotype sample sizes (of non-missing calls) of variant *x* or *y*, respectively:

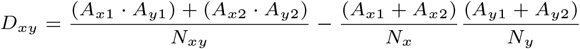

Fundamentally, this relates to the canonical equation of *D*_*xy*_ for a pair of variants *x* and *y* in the following ways:

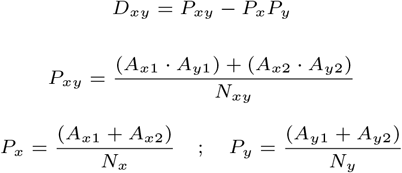

This computation for *D* can be done using a series of matrix calculations, each of which computes different aspects of this calculation:

*P*_*joint*_: N x N matrix where each element at row *x* and column *y* represents *P*_*xy*_ - the probability of joint haplotypes carrying both variants *x* and *y. Ã*_1_ and *Ã*_2_ represent the N x K matrices for each haplotype where NaN elements have been replaced with a 0. The function `is_not_nan` generates a boolean vector where NaN elements are represented by a 0 and non-NaN elements are represented by a 1. The input to is_not_nan is matrix *A*_*geno*_, which represents the sum of the two haplotype blocks *A*_1_ and *A*_2_.

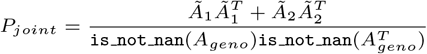

*P*_*mul*_: N x N matrix where each element at row *x* and column *y* represents *P*_*x*_*P*_*y*_ - the product of the probability of the alternate alleles of variant *x* times the same for variant *y* (in genetics terms, the product of the alternate allele fractions of the two variants). *P*_*geno*_ is the variant-wise probability vector where each element is *P*_*x*_ computed by the function `nanmean`, which calculates the mean of an input ignoring NaN values along the axis given. The input to `nanmean` is matrix *A*_*geno*_, which represents *A*_1_ + *A*_2_, and the argument `axis=1` indicates that the variant-wise mean is computed.

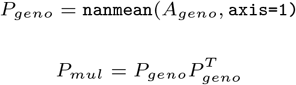

*D*: an N x N matrix where each element at row *x* and column *y* represents *D*_*xy*_ - the coefficient of linkage disequilibrium between variant *x* and variant *y*, as described above.

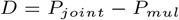

*R*^2^: an N x N matrix where each element at row *x* and column *y* represents 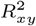 - the phased correlation of variant *x* and variant *y*. Note that this is NOT equal to the square of the Pearson correlation (as discussed in the Unphased section below). We compute the phased *R*^2^ matrix elementwise, and we implement a custom C++ kernel to do so with no memory overhead:

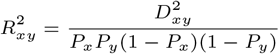

### Computation of the Unphased *R*^2^ LD Matrix

The *R*^2^ LD matrix for unphased data is defined as the square of the correlation matrix *C* for all N variants (“features”) in a dataset, pairwise, with K samples for each variant. *C* is represented by an elementwise calculation, where *A*_*x*_ represents the corresponding row of *A* that represents sample genotypes for each sample in 0/1/2 coding, which can be trivially computed by summing the haplotype blocks *A*_*x*1_ + *A*_*x*2_.

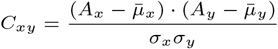

This computation for *C* can be done using a series of matrix calculations where *A*_*geno*_ represents *A*_1_ + *A*_2_, and *µ*_*row*_ and *σ*_*row*_ are column vectors that represent the mean and standard deviation for each variant count in the genotype matrix, respectively:

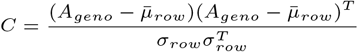

Note that this computation is different from what may be computed by other software for unphased data, such as PLINK, which computes an approximation of the phased *R*^2^ by computing the maximum-likelihood estimate for the haplotype frequencies based on genotype frequencies [12].

### Memory Limitations of Current Hardware

As described above, computing phased LD matrices requires the computation of multiple N x N matrices. For unphased data, one computation of the N x N covariance matrix is required. In both cases, the underlying operations involve large matrix multiplications, which can be highly sped up using GPUs. We have assessed just this step on the CPU and GPU and observed speedups of over 4x on the GPU, even when comparing relatively high-end CPUs to consumer-grade GPUs. However, computing the N x N matrix all at once requires the initialization of *N* ^2^ elements, and for large enough N, would not fit into standard GPU memory. For large biobank-scale datasets, this becomes infeasible, as standard GPU memory cannot accommodate the entire LD matrix at once.

### Scaling LD Computation with ARCHER-LD

In ARCHER-LD, we solve this issue by implementing a chunking approach. Specifically, the genotype matrix (*A*; N x K x 2) is partitioned into discrete subsections to compute chunks of the LD matrix in a parallelized manner. Each subsection *A*_*chunk*_ is a Nchunk x K x 2 matrix, where *N*_*chunk*_ is the number of variants in each chunk. *A*_*chunk*1_ refers to the first chunk and *A*_*chunk*2_ refers to the second chunk. For example, we can compute the block of either the phased or unphased LD matrix that comprises the LD values of variants 1-100 with variants 101-200 (that is, 1::101, 1::102, …, 100::200) at the same time as computing the LD values of variants 1-100 with variants 201-300 on a different GPU.

ARCHER-LD splits up the computation of the entire LD matrix into blocks for each passed *A*_*chunk*_ pair. Noting that the LD matrix is symmetric along the diagonal, we compute only the upper triangle of this matrix for efficiency and chunk the input genotype data in this manner. In all equations above, instances of *A* or *A*_*geno*_ can be replaced with *A*_*chunk*1_ or *A*_*chunk*2_, and any quantities computed from *A* or *A*_*geno*_ such as *µ*_*row*_, *σ*_*row*_, *p*_*geno*_ are similarly replaced by those computed from *A*_*chunk*1_ or *A*_*chunk*2_. This process is visualized in **Figure 1**.

**Fig. 1.**
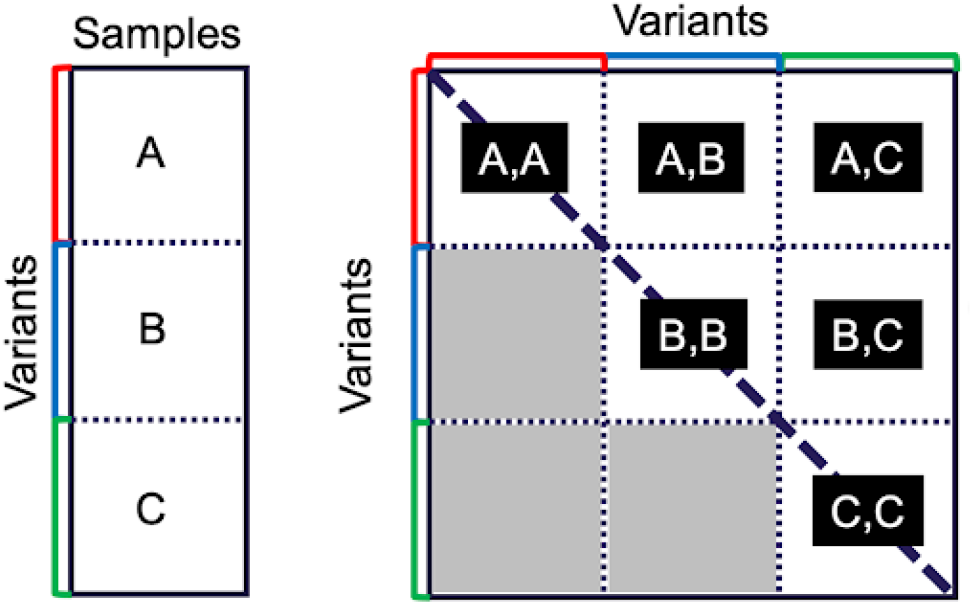
A visualization of the matrix chunking approach (left) and computation of the LD matrix (right). Note that we do not visualize both haplotypes; chunks consist of both haplotypes for phased data or the sum of the haplotypes for unphased data.

We maximize the number of variants that we can load into each chunk by pre-allocating needed contiguous blocks, freeing memory whenever possible, and reusing allocated segments of memory from intermediate computations as frequently as possible, as well as using custom C++ kernels implemented using CuPy. This leads to memory requirements that scale with the number of elements defined as approximately 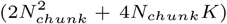 when computing blocks for phased data and 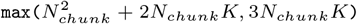 when computing blocks for unphased data.

To implement the chunk-by-chunk calculation across GPUs in multi-node systems, we make use of the Message Passing Interface (MPI) and use CuPy to handle GPU kernel dispatching and operations from within Python. There is a “main” process that schedules chunks to be dispatched to several “compute” processes, where each compute process has one GPU. The “main” process determines the size of the chunks at the beginning based on a user-defined amount of memory available on each GPU. Each “compute” process waits to receive sets of variants and their samples (a subsection of the original N x K x 2 matrix *A*), and upon receiving one, loads the chunked genotype data and performs the computation of the LD matrix (the block) for those sets of variants using basic CuPy operations in a memory-efficient manner, then thresholds and writes the results to disk in the form of a sparse matrix.

### Storage of the Final LD Matrix

To reduce the required storage space and filter out biologically insignificant (i.e. LD values near 0), ARCHER-LD applies a user-defined LD threshold (default: *R*^2^ = 0.1) to compute matrix blocks before writing them to the disk. Notably, the vast majority of variant pairs have close-to-zero LD, and so enforcing even a modest threshold of 0.1 will result in the output matrix being approximately *>*99.999% sparse, which can be stored on disk on the order of gigabytes. We store this final matrix in a Scipy sparse matrix form for ease of access and loading using Python for downstream analyses. There exist tools to load this sparse matrix in R and Stata as well.

### Assessing accuracy by comparing with PLINK2.0

To ensure the accuracy ARCHER-LD’s computations, we compare the results of the phased *R*^2^ computation to the gold standard of PLINK2.0, which provides a function `--r2-phased` to compute the phased *R*^2^ for variants in a given dataset. We performed these computations on chromosome 22 of the phased 1000 Genomes 30x whole-genome-sequencing dataset, comprising 1,066,557 variants and 3202 samples (which includes related individuals). To compare accuracy, we computed the mean squared error, mean absolute error, mean, standard deviation, 99.9th percentile, and absolute maximum of the difference of the results between PLINK2.0 and ARCHER-LD, ignoring non-zero elements to avoid being skewed by the extremely high number of zero values.

We also assess the performance of ARCHER-LD as compared to PLINK in terms of speed, by comparing the amount of walltime that it takes for PLINK to compute the above phased *R*^2^ using a 32-core CPU (an Intel(R) Xeon(R) Gold 6338) with 256GB of RAM on one machine against ARCHER-LD using 8 consumer-grade GPUs (8 NVIDIA GTX 2080TIs) on a different machine.

To assess the scaling of ARCHER-LD with GPU resources and the overall value of using higher-VRAM GPUs, we performed the unphased *R*^2^ computation of ARCHER-LD on a high-performance cluster system (the Department of Energy Argonne Leadership Computing Facility Polaris Supercomputer), where ARCHER-LD had access to more powerful GPUs (512 NVIDIA 40GB A100 GPUs). We did this using all chromosomes of the unphased 1000 Genomes 30x WGS dataset, which has over 120 million variants and over 2500 samples.

## Results

### Accuracy Comparison to PLINK2.0

A comparison of values between PLINK2.0 and ARCHER-LD can be found in **Table 1**. All values except for a single one (owing to precision at the exact boundary of the threshold) were well within floating-point precision and generally in perfect agreement.

**Table 1.** A table of the various metrics used to compare between the *R*^2^ values produced by PLINK2.0 and ARCHER-LD, computed on the difference of (ARCHER-LD - PLINK2.0)

| Metric | Value |
| --- | --- |
| Mean Squared Error | $2.29 \times 10^{-9}$ |
| Mean Absolute Error | $2.71 \times 10^{-7}$ |
| 99.9th%ile of Absolute Differences | $8.56 \times 10^{-7}$ |
| Absolute maximum | 0.1<sup>a</sup> |

<sup>a</sup>This value occurred from a single variant. ARCHER-LD provided a value of 0.1 (to 6 decimal places) and PLINK2.0 provided 0.0, likely due to tiny differences in precision before thresholding. The largest absolute difference disregarding this single variant is 6.22e-06.

### Speed Comparison to PLINK2.0

PLINK2.0 used 128GB of RAM and 64 threads per its default settings (which uses half of the memory available) from the resources provided to it. The overall run time to compute all phased pairwise LD on chromosome 22 for PLINK2.0 was 81 minutes and 18 seconds, excluding time taken to load and convert the dataset (preconverted to PLINK binary files from VCF using PLINK). ARCHER-LD computed all of the phased pairwise LD on chromosome 22 in approximately 8 minutes and 17 seconds after taking 1 minutes and 33 seconds to convert the chromosome 22 VCF to Zarr format, for a total of 9 minutes and 50 seconds using 8 NVIDIA GTX 2080TI GPUs that each had 12GB of VRAM, a speedup of approximately 8 times over PLINK.

### Scaling up to hundreds of GPUs for genome-wide LD

When scaling up to 1000 Genomes, we computed on the aforementioned 30x WGS 1000 Genomes dataset (≈120 million variants and ≈2500 samples) the entire genome-wide pairwise unphased LD matrix (*>*1e16, or 10 quadrillion elements) in under 6 hours using 512 NVIDIA 40GB A100 GPUs on the DOE ALCF Polaris supercomputer, representing a major breakthrough in our ability to compute genome-wide linkage disequilibrium.

## Conclusion

ARCHER-LD is written in Python using CuPy and custom C++ kernels and is publicly available. ARCHER-LD allows researchers to compute genome-wide linkage disequilibrium matrices at biobank scale and allows them to design experiments that leverage the full extent of their genomic data through the use of long-range LD and without the need for external reference panels. ARCHER-LD will enable the design of more accurate and population-specific analyses, particularly for studies in historically underrepresented populations in existing databases.

## Author contributions statement

RK wrote the code for this manuscript and the initial draft of this manuscript. RK and AV conceived of the idea for this project. RK, PS, DZ, CC, MC, BFV, MDR, and AV all contributed to the theoretical underpinnings of the approach. AR, TNN, MT, and RKM all provided guidance and support for the technical implementation and testing. All authors revised the manuscript and approved the final draft of the manuscript.

### Acknowledgments

RK was supported by the National Human Genome Research Institute of the National Institutes of Health (T32HG000046). The funders played no role in study design, data collection, analysis and interpretation of data, or the writing of this manuscript.

## Code availability

The code and documentation for ARCHER-LD is available at https://github.com/rachitk/ARCHER-LD.

A container is available to run ARCHER-LD, and we will also deposit the version of the code at publication into a data-sharing repository to serve as a permanent identifier.

## Supplementary Analyses

### Speed comparison in the Penn Medicine Biobank

To further assess how well ARCHER-LD scales in larger datasets, we used the Penn Medicine Biobank (PMBB) dataset with 57170 samples. We compared the speed of computing the unphased LD on chromosomes 1 and 22 separately using PLINK2.0 and ARCHER-LD. It is important to note that this is not a true 1:1 comparison, as PLINK computes the maximum-likelihood estimate (MLE) of the LD statistic for unphased data rather than the true statistic. Computing the MLE is generally less computationally expensive than computing the true LD, which is why it is done in large, unphased datasets.

For these tests, PLINK2.0 had access to 256GB of RAM and 64 threads on an AMD Genoa EPYC 9374 CPU while ARCHER-LD had access to 8 NVIDIA Blackwell B200 GPUs (each with 180 GB of VRAM, though of note, we only made use of approximately 30GB of VRAM per GPU in these tests due to cluster restrictions at the time of testing). The results can be seen in **Table 2**.

**Table 2.** Speed of ARCHER-LD and PLINK in computing unphased LD on the PMBB dataset)

| Chromosome | Variants | PLINK2.0 | ARCHER-LD | Speedup |
| --- | --- | --- | --- | --- |
| 1 | 1,380,131 | 22:40:28 | 00:49:13 | $\approx 27.6$ |
| 22 | 326,449 | 01:41:05 | 00:05:15 | $\approx 19.2$ |

As can be seen, ARCHER-LD’s speedup compared to PLINK appears to grow with dataset size, with its speedup on unphased data appearing to be quite significant.

### Variant chunk sizes with GPU memory and sample sizes

We plot how the chunk sizes that can be computed on a given GPU for a given dataset vary with GPU memory (VRAM) and dataset sample size in **Figure 2** and **Figure 3**, respectively.

**Fig. 2.**
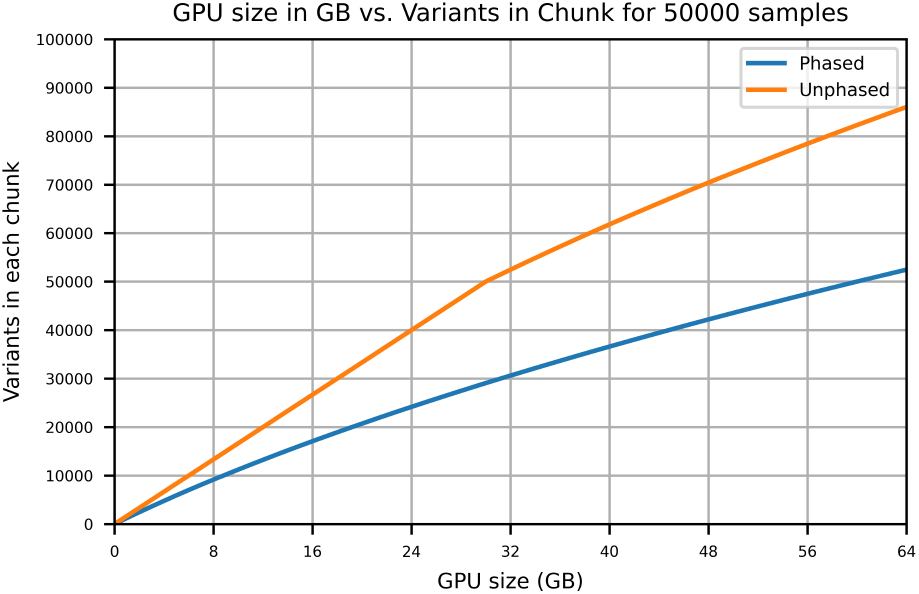
A visualization of variant chunk sizes for different GPU memory sizes from 0GB to 64GB for an arbitrary fixed sample size (50000). Note the discontinuity when the variant chunk size equals the number of samples at 50000 for unphased data, representing when the size of the variant matrices exceeds the size of the LD matrix block computed.

**Fig. 3.**
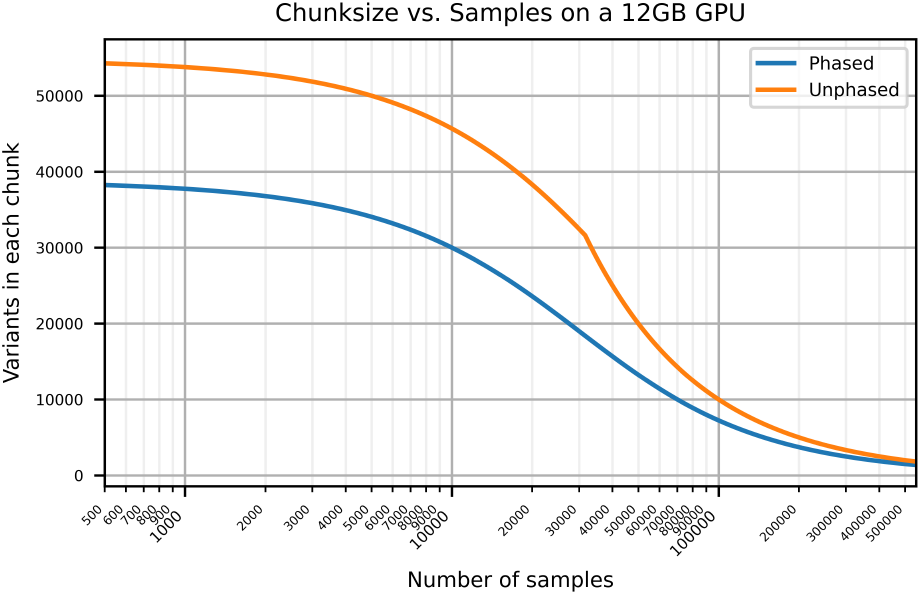
A visualization of variant chunk sizes for different sample sizes from 500 to 500,000 for a fixed GPU memory (12 GB of VRAM, typical of many modern consumer-level GPUs). Note the discontinuity when the variant chunk size equals the number of samples at approximately 30000 for unphased data, representing when the size of the variant matrices exceeds the size of the LD matrix block computed.

## References

1. Montgomery Slatkin. Linkage disequilibrium — understanding the evolutionary past and mapping the medical future. Nature reviews. Genetics, 9(6):477–485, June 2008.

2. Alkes L. Price, Michael E. Weale, Nick Patterson, Simon R. Myers, Anna C. Need, Kevin V. Shianna, Dongliang Ge, Jerome I. Rotter, Esther Torres, Kent D. Taylor, David B. Goldstein, and David Reich. Long-range LD can confound genome scans in admixed populations. American Journal of Human Genetics, 83(1):132–135; author reply 135–139, July 2008.

3. Effie W. Petersdorf and Colm S. O’hUigin. The MHC in the Era of Next-Generation Sequencing: Implications for Bridging Structure with Function. Human immunology, 80(1):67–78, January 2019.

4. Evan Koch, Mickey Ristroph, and Mark Kirkpatrick. Long Range Linkage Disequilibrium across the Human Genome. PLoS ONE, 8(12):e80754, December 2013.

5. Pankhuri Singhal, Yogasudha Veturi, Scott M. Dudek, Anastasia Lucas, Alex Frase, Kristel van Steen, Steven J. Schrodi, David Fasel, Chunhua Weng, Rion Pendergrass, Daniel J. Schaid, Iftikhar J. Kullo, Ozan Dikilitas, Patrick M.A. Sleiman, Hakon Hakonarson, Jason H. Moore, Scott M. Williams, Marylyn D. Ritchie, and Shefali S. Verma. Evidence of epistasis in regions of long-range linkage disequilibrium across five complex diseases in the UK Biobank and eMERGE datasets. American Journal of Human Genetics, 110(4):575–591, April 2023.

6. Christian Benner, Aki S. Havulinna, Marjo-Riitta Jarvelin, Veikko Salomaa, Samuli Ripatti, and Matti Pirinen. Prospects of Fine-Mapping Trait-Associated Genomic Regions by Using Summary Statistics from Genomewide Association Studies. American Journal of Human Genetics, 101(4):539–551, October 2017.

7. Yuxin Zou, Peter Carbonetto, Gao Wang, and Matthew Stephens. Fine-mapping from summary data with the “Sum of Single Effects” model. PLOS Genetics, 18(7):e1010299, July 2022.

8. Brent S Pedersen and Aaron R Quinlan. Cyvcf2: Fast, flexible variant analysis with Python. Bioinformatics, 33(12):1867–1869, June 2017.

9. Fsspec/kerchunk. python filesystem spec, May 2026.

10. Eric Czech, Will Tyler, Tom White, Ben Jeffery, Timothy R Millar, Benjamin Elsworth, Jeremy Guez, Jonny Hancox, Konrad J Karczewski, Alistair Miles, Sam Tallman, Per Unneberg, Rafal Wojdyla, Shadi Zabad, Jeff Hammerbacher, and Jerome Kelleher. Analysis-ready VCF at Biobank scale using Zarr. GigaScience, 14:giaf049, January 2025.

11. Josh Moore and Susanne Kunis. Zarr: A Cloud-Optimized Storage for Interactive Access of Large Arrays. Proceedings of the Conference on Research Data Infrastructure, 1, September 2023.

12. Christopher C Chang, Carson C Chow, Laurent CAM Tellier, Shashaank Vattikuti, Shaun M Purcell, and James J Lee. Second-generation PLINK: Rising to the challenge of larger and richer datasets. GigaScience, 4(1):s13742–015–0047–8, December 2015.

